# *In vitro* community synergy between bacterial soil isolates can be facilitated by pH stabilisation of the environment

**DOI:** 10.1101/347039

**Authors:** Jakob Herschend, Klaus Koren, Henriette L. Røder, Asker Brejnrod, Michael Kühl, Mette Burmølle

**Author notes:** These authors contributed equally to this study (Jakob Herschend, Klaus Koren). **Correspondence:** Mette Burmølle, Universitetsparken 15 Bldg. 1, 2100 Ø, Denmark.

## Abstract

Composition and development of naturally occurring microbial communities is defined by a complex interplay between the community and the surrounding environment and by interactions between community members. Intriguingly, these interactions can in some cases cause community synergies where the community is able to outperform it single species constituents. However, the underlying mechanisms driving community interactions are often unknown and difficult to identify due to high community complexity. Here we show how pH stabilisation of the environment through the metabolic activity of specific community members acts as a positive inter-species interaction driving *in vitro* community synergy in a model consortium of four co-isolated soil bacteria: *Microbacterium oxydans*, *Xanthomonas retroflexus*, *Stenotrophomonas rhizophila* and *Paenibacillus amylolyticus*. Using micro-sensor pH measurements to show how individual species change the local pH micro-environment, and how co-cultivation leads to a stabilised pH regime over time. Specifically, *in vitro* acid production from *Paenibacillus amylolyticus* and alkali production primarily from *Xanthomonas retroflexus* lead to an overall pH stabilisation of the local environment over time, which in turn resulted in enhanced community growth. This specific type of interspecies interaction was found to be highly dependent on media type and media concentration, however similar pH drift from the individual species could be observed across media variants.

**Importance:** We show that *in vitro* metabolic activity of individual members of a synthetic, co- isolated model community presenting community synergistic growth arises through the inter-species interaction of pH stabilization of the community micro-environment. The observed inter-species interaction is highly media specific and most pronounced under high nutrient availability. This adds to the growing diversity of identified community interactions leading to enhanced community growth.

## Introduction

Microbial communities are ubiquitous in natural and man-made environments and are routinely being applied for e.g. crop-management (1), bioremediation (2), waste-water treatment (3) and bio-energy production (4, 5). Hence, in terms of biotechnological applicability and environmental ecology, understanding key factors affecting microbial community development is indispensable (6). The actively growing community in a natural habitat is predominantly defined in diversity and composition by environmental factors e.g. O2, pH, salinity and temperature (7–11), where the chemical micro-environment is characterized by steep chemical gradients susceptible to rapid changes. By example, pH is recognized as an important factor for species composition in e.g. soil (11–13), as different species prefer specific pH regimes (14, 15). Albeit the strong environmental effect, microbial interactions also influence community composition e.g. through molecular mechanisms such as cooperative cross-feeding (16–18) and cross-protection from anti-biotics (19, 20) or through competition by toxin secretion (21). An additional mode of interaction is based on bacteria’s ability to alter their local environment, e.g. by changing the local pH micro-environment by consumption of specific resources, secretion of metabolites or through the bio-chemical processes from metabolic activity causing a proton turnover (22, 23). Microbial pH drift of the environment is well known from several types of host-associated environments such as e.g. the human-associated vaginal (24) and oral (25) microbiomes, and from the well-known syntrophic relationship of industrial yoghurt production by *Lactobacillus bulgaricus* and *Streptococcus thermophiles* (26–29). Recently Ratzke *et al.* (2018) showed through *in vitro* studies that in unique cases bacteria may even cause pH drift to such an extent that it becomes detrimental for the population, a phenomenon termed ecological suicide (15). As pH is an important parameter for microbial life, changing the pH environment will affect both the microbial population causing the change and neighbouring community members; Such pH interactions in co-cultures and how their outcome can be modelled have been elegantly documented in detail for *in vitro* co-cultures by Ratzke and Gore (2018) (14). Using specific laboratory isolates Ratzke and Gore (2018) showed that the outcome of pH driven interactions can be modelled when the pH drift and pH growth optimum is known for the interaction partners. The outcome of the interaction could then be categorized as e.g. bi-stability, successive growth, extended suicide or stabilization of growth. By example, stabilization defines a scenario where two bacteria, which on their own would create a detrimental pH environment, can co-exist by canceling each other’s pH drift of the environment.

The range of interactions occurring in bacterial communities often facilitates the emergence of properties, which are only observed in a community setting and not from its single species members, referred to as community-intrinsic properties (30). An example of a community-intrinsic property is the synergistic biofilm formation recorded by Ren et al. (2015) (31) for a model community consisting of four co-isolated soil bacteria; *Stenotrophomonas rhizophila*, *Xanthomonas retroflexus*, *Microbacterium oxydans* and *Paenibacillus amylolyticus.* Work on this community has established that co-cultivation leads to enhanced biofilm formation, that all four species increase in cell counts through biofilm co-cultivation and that all four species are indispensable for the synergy to occur (31). Later studies has hinted that the synergy can be linked to a specific spatial organisation of community members during co-cultivation in biofilms (32), and meta-transcriptomics (33) and - proteomics (34) studies have identified amino acid cross-feeding as a potential driver for the synergy. However, the impact of the community on its surrounding environment, and the mutual community-environment interplay, has not been explored. Hence in the framework of inter-species interactions occurring through changing the local environment, we here applied high resolution microsensor measurements of pH and O_2_ (35, 36) in liquid cultures and solid surfaces to elucidate the role of the chemical micro-environment on the observed community synergy from this model community. In line with observations from Ratzke and Gore (2018), we find that three community members individually drive pH to un-favourable conditions hampering their own growth, whereas co-cultivation leads to a stabilisation of the environment, promoting community synergy.

## Results

### Bacterial interactions on an agar plate

The species were spotted in pairs of two on agar plates (50% TSA with congo red and coomassie brilliant blue G250), to screen for interactions between the species. Interactions would be detected by visual changes in colony morphology. After two days of incubation, colony morphology of *Paenibacillus* changed when spotted against *Xanthomonas*, *Stenotrophomonas* or *Microbacterium* (Fig. 1), compared to when spotted against itself. The changed *Paenibacillus* colony was increasingly red, indicating enhanced binding of congo red, and the colony texture was disordered in the peripheral part opposing the other species. The visually disordered part displayed directional growth towards the opposing colony, indicating attraction. The reaction was strongest against *Xanthomonas.* No visible interactions were observed among the other species pairs, as judged by colony morphology (Fig. S1).

**Figure 1:**
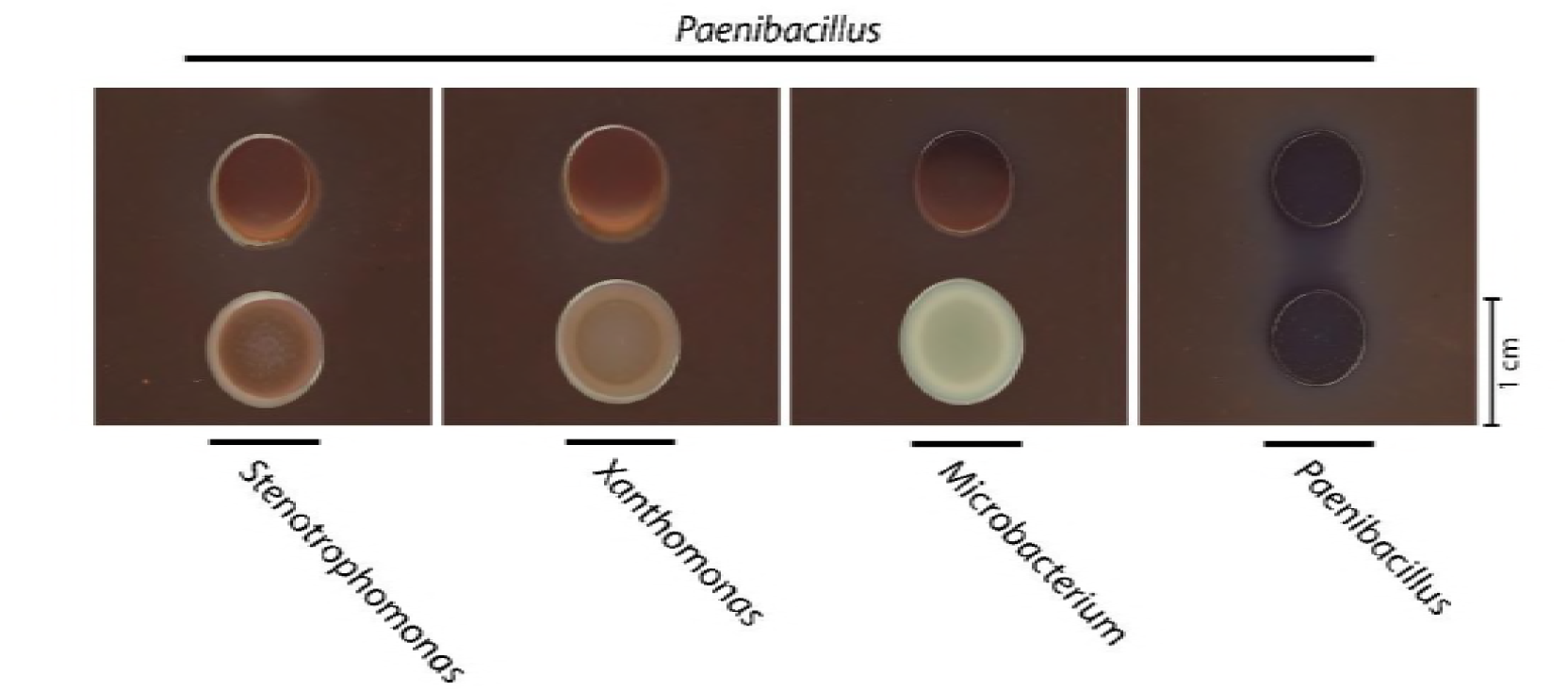
Two-species interactions with *Paenibacillus*. *Paenibacillus* colony morphologically changed when spotted close to *Stenotrophomonas*, *Xanthomonas* and *Microbacterium* colonies on congo red plates. The part of the *Paenibacillus* colony opposing the other species turned light red and grew directionally towards the opposing species. No morphological changes occurred when *Paenibacillus* was spotted against itself.

### The chemical micro-environment in the agar

As the morphological change of *Paenibacillus* indicated directional growth, it was hypothesized that *Xanthomonas* (and to a lesser extend *Stenotrophomonas* and *Microbacterium*) modified the chemical environment in the agar causing attraction of *Paenibacillus*. To probe the chemical micro-environment of the interaction zone between the colonies on agar plates, the zone was mapped in a 2.5 × 2.5 mm grid-structure using O_2_ and pH micro-sensors mounted on a 3D motorised micro-manipulator (Fig. 2). The experimental setup is presented in Fig. S2. A visible morphological change of the *Paenibacillus* colony occurred from day one to two. According to the pH map (Fig. 2), the pH changed after one and two days of incubation. After one day of incubation, pH increased in the area around *Xanthomonas* and *Stenotrophomonas*, as compared to the pH of 50% TSA medium-based agar (indicated by black arrow, Fig 2). Simultaneously, the pH in parts of the *Paenibacillus* colony periphery not facing the interaction zone decreased to ~ pH 6.5. No change in pH was observed close to *Microbacterium* colonies. After two days of incubation, the pH in the interaction zone increased to pH >8. At the periphery of the *Paenibacillus* colony opposite the interaction zone, the pH was still below pH 8. The pH data showed that *Xanthomonas* and *Stenotrophomonas* alkalized the environment when cultured with TSB as the nutrient source, whereas *Paenibacillus* acidified its environment. No clear trend was observed for *Microbacterium*.

**Figure 2:**
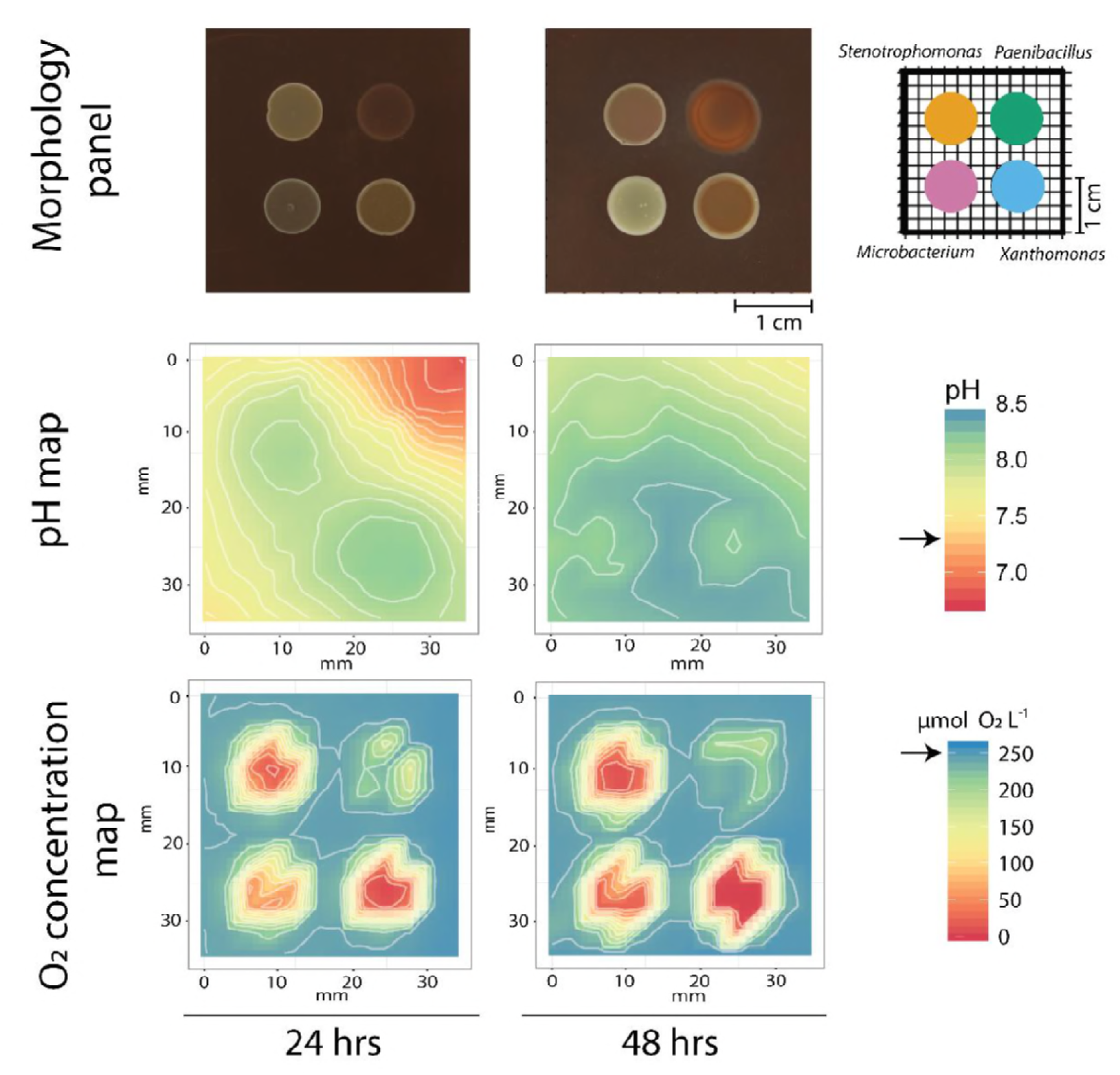
Mapping of O_2_ and pH in the interaction zones of *Xanthomonas*, *Stenotrophomonas*, *Microbacterium* and *Paenibacillus* grown on 50% TSA plates. Arrows on legend bars indicate pH and O_2_ concentration in 50% TSA agar without bacteria. The pH and O_2_ concentrations were measured 100 µm below the surface of the agar at each 2.5 × 2.5 mm grid position. Morphology panels show the interaction of *Paenibacillus* occurring on congo red plates and the positioning of the individual species on the plate. Panels with pH measurements at 24 hours show increased pH around *Stenotrophomonas* and *Xanthomonas* colonies, with an alkalization of the media towards pH 8. In the periphery of the *Paenibacillus* colony opposite the interaction zone, the agar was acidified towards pH 6.5. After two days of growth of *Xanthomonas* and *Stenotrophomonas*, the pH in the majority of the interaction zone was enhanced to pH ≥8.0. After 24 and 48 hrs growth, *Xanthomonas*, *Stenotrophomonas* and *Microbacterium* deprived the agar of O_2_, leaving the respective colony centres anoxic. Only a small O_2_ depletion was measured in the periphery of the *Paenibacillus* colony.

The O_2_ concentration map indicated strong O_2_ depletion by *Stenotrophomonas*, *Xanthomonas* and *Microbacterium* after one and two days of incubation, where the central parts of these colonies reached anoxia. In contrast, only a weak O_2_ depletion was recorded near the periphery of the *Paenibacillus* colony, after both one and two days of incubation.

### Measurement of pH and growth in liquid co-cultures

With the opposing trend of pH drift from *Stenotrophomonas* and *Xanthomonas* compared to *Paenibacillus,* it was hypothesised that environmental pH stabilisation, similar to that observed by Gore and colleagues (14, 15), could also be a driver for the observed community synergy observed by Ren et al. (31) and Hansen et al. (33) in static liquid cultures. Mono-, dual- and four-species cultures were grown in 24-well polystyrene plates with measured endpoint pH and individual counts of colony forming units (CFU) from all species. Oppositely to Ren et al. (31) who specifically quantified the biofilm constituents (bacterial biofilm cells and biofilm matrix), the present study quantified cell content in the entire well, as selective pH measurements in the biofilm fraction of 24 well plates were impractical. In line with the observations by Ren et al. (31), *Xanthomonas* was the most abundant species in the four-species community contributing >95% of the total cell counts (Fig. 3a). The four-species consortium yielded higher total CFU counts than the best single species culture, e.g. *Xanthomonas*, and counts equalled the sum of single species, indicating some level community synergy (Fig. 3b). Cell counts of *Xanthomonas* and *Paenibacillus* were higher in the four-species consortium as compared to their respective mono-cultures. Oppositely, cell counts of *Stenotrophomonas* and *Microbacterium* were reduced when included in the four-species consortium (Fig. 3c). Similar to the observations from agar plates, *Xanthomonas* and *Stenotrophomonas* alkalised the medium when cultured individually in TSB, driving pH ≥ 8, whereas *Paenibacillus* produced acid, driving pH < 6 (Fig 3d, e and Fig. S3). In contrast to the observations from agar plates, a slight acidification by *Microbacterium* was detected in static liquid TSB cultures (Fig. S3).

**Figure 3:**
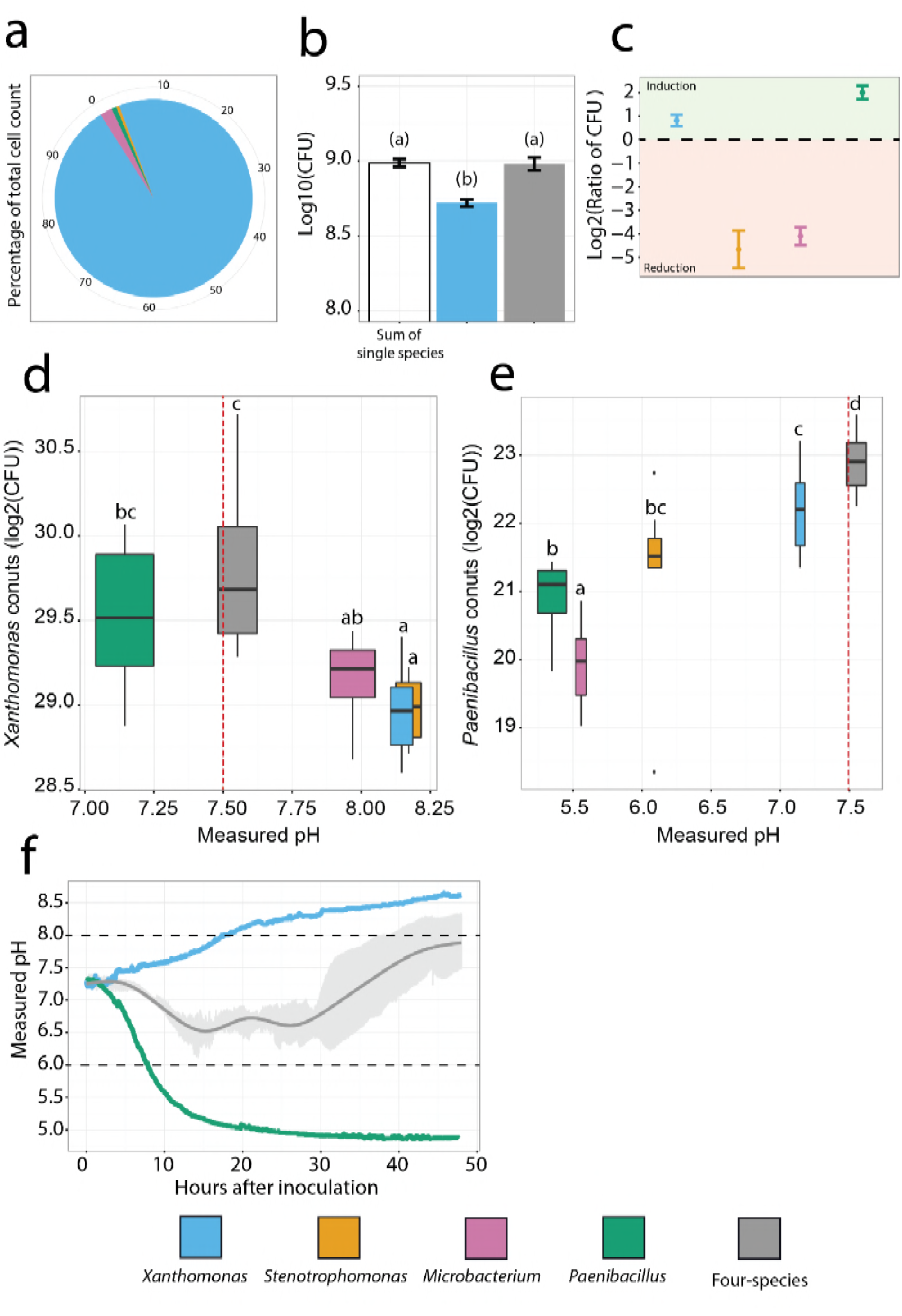
Growth and composition of the four-species community, along with its effect on the local environment. a) Species distribution based on CFU counts in the four-species community, with *Xanthomonas* composing >95% of the four-species community (n = 10 biological replicates). b) Community productivity by total cell counts (log10(CFU) of the four-species community, compared to best single species (*Xanthomonas*) and the sum of single species. Bars indicate standard error and dissimilar letters indicate significant differences with p<0.05 (GLM) (n = 10 biological replicates). c) Species dynamics in the four-species consortium, compared to single species populations. Bars indicate standard error. Cell counts of both *Xanthomonas* and *Paenibacillus* were higher when co-cultured in the four-species consortium, whereas cell counts of *Microbacterium* and *Stenotrophomonas* were reduced (n = 10 biological replicates). d+e) CFU counts of *Xanthomonas* and *Paenibacillus* respectively, in mono- and co-cultures, mapped with endpoint pH for each culture after 48 hrs of incubation. Spearman’s ranked correlation between CFU and endpoint pH. Statistical grouping of CFU, with dissimilar letters, e.g. a and b, indicating significant differences with p<0.0.5 (GLM) (n = 10 biological replicates). Box width represents two times the standard error of the measured endpoint pH in each culture. Co-cultures are labelled with the colours of the included species. Counts of the four-species consortia are labelled in grey. Red dotted line represents pH in the media without inoculation. f) Time trace of measured pH during growth of *Xanthomonas*, *Paenibacillus* and the four-species consortium. Data from *Paenibacillus* and *Xanthomonas* represents a single biological replicate. Additional replicates were made to verify the single-species trend, but these are not included in the data representation. Data from the four-species consortium represent the smoothed average of three biological replicates (dark grey line). Standard deviation for each measured time point is plotted as bars (light grey). The dotted lines mark optimal pH growth range for each of the four species, as estimated by growth on buffer stabilized 50% TSA plates (supplementary fig. 4).

When comparing end-point pH and CFU counts of individual species in mono-, dual- and four-species cultures, it was apparent that different species compositions resulted in unique end-point pH and CFU counts for each culture, as seen by e.g. endpoint pH and CFU counts of *Xanthomonas* or *Paenibacillus* (Fig 3d, e, respectively). For *Xanthomonas*, the mono-culture or co-cultivation with either *Stenotrophomonas* or *Microbacterium* resulted in endpoint pH ≥ 8 and lower CFU counts of *Xanthomonas*, compared to cultures including the acid-producer *Paenibacillus*. Co-cultivation of *Xanthomonas* and *Paenibacillus* or as part of the four-species consortium resulted in significantly higher CFU and lower endpoint pH. In support, a Spearman’s ranked correlation showed a significant (p-value = 0.0058) weak negative correlation (ϱ_Spearman_ = −0.38) between pH and CFU, indicating that higher pH lead to reduced CFU counts (Fig. 3d).

For *Paenibacillus* (Fig. 3e), an opposite trend was observed as co-cultivation with the strong alkali-producers, e.g. *Stenotrophomonas* or *Xanthomonas,* yielded higher CFU counts, and co-cultivation in the four-species consortium resulted in the significantly highest *Paenibacillus* CFU counts. A strong positive (ϱ_Spearman_ = 0.82) and significant (p-value < 0.0001) Spearman’s ranked correlation indicated a positive relationship between endpoint pH and CFU. While co-cultivation of *Xanthomonas* or *Paenibacillus* with other species or each other generally resulted in increased CFU counts, co-cultivation of *Stenotrophomonas* or *Microbacterium* with other species generally affected the growth of *Stenotrophomonas* or *Microbacterium* negatively (Fig. S3a, b). Hence, other factors than pH are also important for the community growth. No statistically significant Spearman’s correlation was found between pH and CFU counts for *Microbacterium* and *Stenotrophomonas*.

### Stabilisation of pH over time

Measurements of endpoint pH and CFU showed a general trend that co-cultivation of *Paenibacillus* or/and *Xanthomonas* stabilized pH between the observed extremes of their respective mono-cultures, while simultaneously yielding increased CFU counts. To verify that the pH stabilisation occurred throughout the cultivation period and was not just an endpoint artefact, pH was measured over time in mono-cultures of *Xanthomonas*, *Paenibacillus* and the four-species culture with measurements every 5 minutes over 48 hrs. In *Xanthomonas* mono-cultures pH was raised to above 8 within the first day, while *Paenibacillus* mono-cultures acidified the environment to pH 5 within the same time frame. In contrast, growth of the four-species consortium stabilised pH between pH 6 and 8 (Fig. 3f). To evaluate the optimal pH growth range of the individual species, each species was spotted onto pH stabilized 50% TSA plates (Fig. S4). *Stenotrophomonas* and *Xanthomonas* were able to grow at pH 6-8, with no visible growth below pH 6 and above pH 8. *Microbacterium* and *Paenibacillus* grew well between pH 6 and 8, with reduced growth between pH 8 and 9. No growth was recorded for *Microbacterium* or *Paenibacillus* below pH 6.

As *Xanthomonas* and *Paenibacillus* were also able to mutually enhance each other’s growth in dual-culture, pH was continuously measured in this dual-culture to verify that they would also cause pH stabilisation over time. The pH stabilisation was indeed observed, indicating that at least part of pH stabilisation was facilitated through the growth of these two species together (Fig. S5). Complementary measurements showed that the pH stabilisation occurred simultaneously throughout the entire well, as no spatial pH gradients were found between top and bottom (data not shown).

As tryptic soy broth (TSB) is rich in peptides, the observed pH increase for *Xanthomonas* and *Stenotrophomonas* cultures was believed to be caused by the release of ammonia from peptide degradation. Ammonia production was quantified by performing endpoint ammonium measurements after two days of growth, as proton uptake by ammonia would lead to formation of ammonium (Supplementary section on “Nitrogen flux and its impact on community composition”, specifically Table S1 and Fig. S6). Mono-cultures of *Xanthomonas* and *Stenotrophomonas* contained significantly higher concentrations of ammonium (p-value < 0.05, Table S1 and Fig. S6) than those found in TSB, indicating that the change in pH was caused by active degradation of amino acids and release of ammonia. A significantly higher concentration of ammonium was also measured for the four-species consortium (Table S1 and Fig. S6).

We expect the observed pH decrease in *Paenibacillus* cultures to be the result of a fermentative metabolism. When tested by Hugh-Leifson test *Paenibacillus* shows acids production under anaerobic conditions supporting its ability to perform fermentation (data not shown), additionally genome analysis has revealed the genomic potential for lactate production (data not shown). As a potential fermentative metabolism would be favoured in an anoxic environment, oxygen profiles were made on the 24 wells plates over time. Oxygen profiles showed that oxygen was depleted within the first 1 hrs of inoculation for all single- and the four-species cultures in 24 well plates (data not shown). As the environment turned anoxic, growth of the non-fermenting species *Xanthomonas*, *Stenotrophomonas* and *Microbacterium*, would have to rely on alternative electron acceptors. Nitrogen flux of nitrate, nitrite and nitrous oxide was measured in the cultures, and has been summarized in Table S1 and Fig. S6. In short, *Xanthomonas, Paenibacillus* and *Microbacterium* were found to respire on nitrate, which was believed to allow continued growth of these species after oxygen depletion. A more detailed description of nitrogen flux and a complementary genome analysis of each species is available in the supplementary text section “Nitrogen flux and its impact on community composition”. We speculate that *Stenotrophomonas* was the least fit to thrive in the community, as it lacked the ability to perform anaerobic dissimilatory nitrate respiration, as compared to the other species.

### Stability of the pH related interaction

The observed pH related interaction resembled the phenomenon reported by Gore and colleagues (14, 15) where bacteria with opposite pH drive can stabilise each other’s growth, referred to as stabilization. To address the stability of the pH related interaction, counts of individual species and end-point pH were collected for mono and co-cultures in varying strength of TSB and with M9 and LB media as alternative nutrient sources. LB was included due to its complexity, to address if the pH synergy would prevail in other types of complex media. M9 was made with 0.5% tryptone and 0.5% glucose to have a defined and simple growth medium. CFUs and pH were assessed for mono- and four-species cultures after both 24 and 48 hrs of incubation, whereas CFUs and pH for dual-species combinations were only assessed at 48 hrs. Glucose concentrations found in the tested media are within the range of carbohydrates in soil (0.1% (37) to 10% (38)), with M9 having 0.5% and TSB 0.25% glucose. Across media variants and time points, *Xanthomonas* was generally among the species with the highest single species counts (Fig. S9) and in the four-species community it was consistently the most abundant member (Fig. S10). *Xanthomonas* (Fig. 4a, b and Fig. S11) and *Stenotrophomonas* (Fig. S12) increased pH in all tested media after 48 hrs, while *Paenibacillus* (Fig. 4c, d and Fig. S13) caused acidification in both TSB and LB, but not in M9. *Microbacterium* (Fig. S14) slightly acidified TSB based media and increased the pH in LB medium. No clear trend was observed in M9 for *Microbacterium*. The observed synergy from full strength TSB was found to cease with decreasing TSB concentrations, with summed CFU counts of the four-species community in 50% and 20% TSB not being significantly higher than counts of the best single species. At 48 hrs and in 50% TSB the four-species community still had higher average counts than the best single species, indicating that the synergistic interaction to some extent was still in play (Fig. S15). The CFU based synergy was detected in LB medium at 24 hrs, but not at 48 hrs. No synergy was observed when the four-species community was grown in M9 media. Hence, the synergy seemed highly media and concentration dependent.

**Figure 4:**
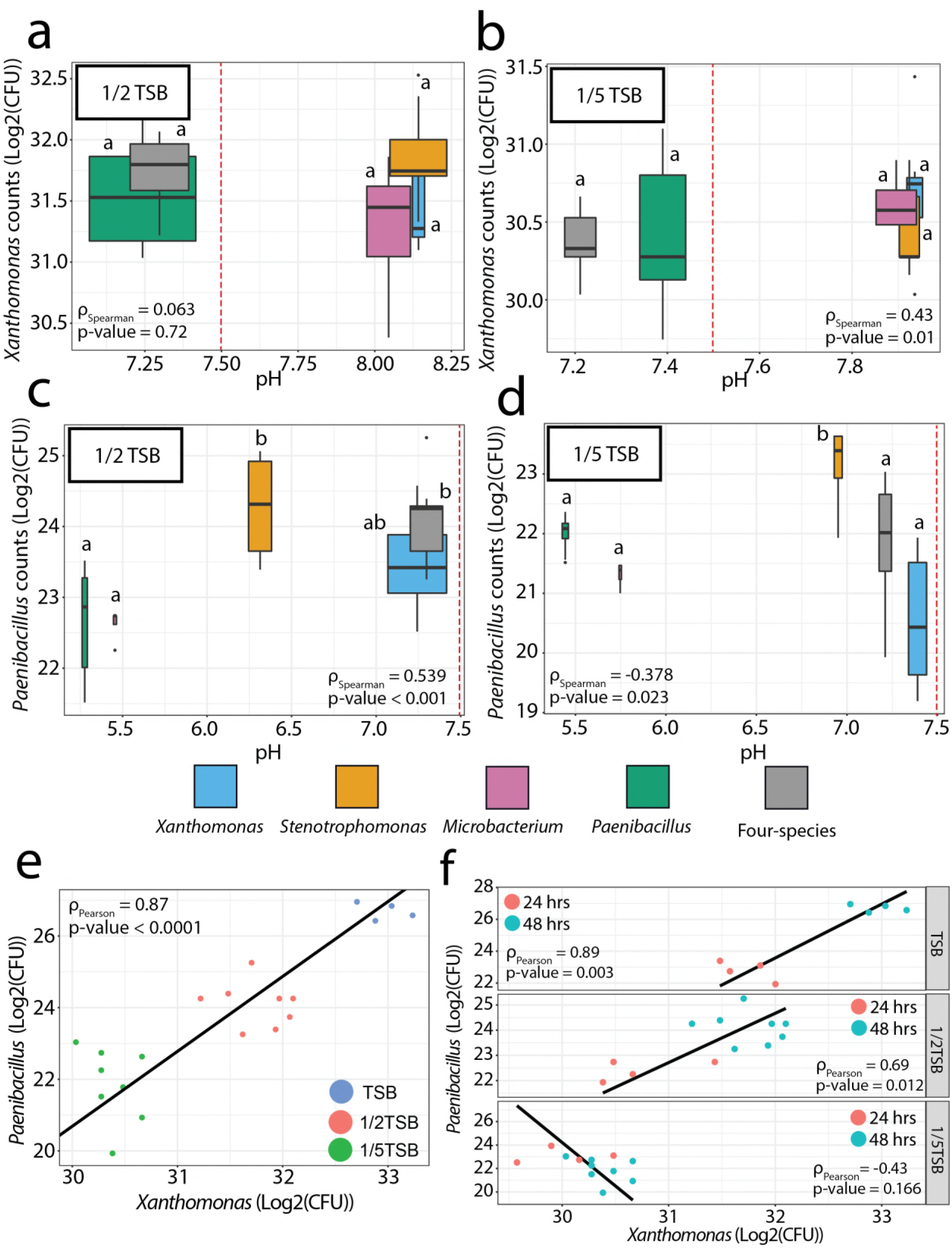
Growth of *Xanthomonas* and *Paenibacillus* across tested media concentrations. a-d) CFU counts of *Xanthomonas* (a-b) and *Paenibacillus* (c-d) respectively, in mono- and co-cultures with 50 and 20% TSB plotted against endpoint pH for each culture after 48 hrs of incubation (n = 8 biological replicates, at 48 hrs of growth). Spearman’s correlation between CFU and end point pH are presented in each panel. Statistical grouping of CFU is presented by dissimilar letters indicating significant differences with p < 0.05 (GLM). Box width represents two times the standard error of the measured endpoint pH in each culture. Co-cultures are labelled with the colours of the included species and the counts from the four-species community is labelled grey. Red dotted line represents pH in the media without inoculation. The positive effect of pH stabilisation on *Xanthomonas* disappear with decreased media concentration, whereas the effect is still present for *Paenibacillus* at 50% TSB. e) Counts of *Xanthomonas* and *Paenibacillus* across variants of TSB when cultured as part of the four-species community. A strong positive and significant Pearson’s correlation between counts of both species (log2(CFU)) indicates that these two species responds to each other’s growth across media concentrations. (n = 4-8 biological replicates, measured at 48 hrs) f) Counts of *Xanthomonas* and *Paenibacillus* when cultured as part of the four-species community across variants of TSB and across time points. Pearson’s correlation between counts of both species (log2(CFU)) (n = 4-8 biological replicates, with n = 4 at 24 hrs and n = 8 at 48 hrs). *Xanthomonas* and *Paenibacillus* has a positive effect on each other in full strength and 50% TSB, while also following each other’s growth.

As *Xanthomonas* and *Paenibacillus* were hypothesised to be the main drivers behind the observed pH interaction mediating the synergy, analysis of end-point pH and CFU across co-cultures and media for these two species was applied to unravel when the pH related effect disappeared with decreasing media concentration. For *Xanthomonas* (Fig. 4a-b) the positive effect observed during co-cultivation in full strength TSB disappeared in 50% and 20% TSB, yielding comparable CFU counts for mono- and co-cultures. Hence, the pH-related effect ceased with decreasing media concentration for *Xanthomonas*. For *Paenibacillus* (Fig. 4c-d) a significant Spearman’s ranked correlation (ϱ_Spearman_ = 0.539, p-value < 0.001) was found between CFUs and pH in 50% TSB, with co-cultures of e.g. *Stenotrophomonas* and the four-species culture also providing significantly higher CFU counts. Combined, this indicated a continued relationship between pH stabilisation and increased CFUs. For 20% TSB the pH-mediated interaction also ceased for *Paenibacillus*, as co-cultivation yielded comparable CFU counts to that of mono -cultures.

*Xanthomonas* and *Paenibacillus* was found to be the main drivers for the community synergy from full strength TSB having a pH stabilizing interaction. However, the pH related interaction between *Xanthomonas* and *Paenibacillus* was observed to be both media and concentration dependent, with high nutrient loadings being required for the interaction to occur. In support of an interaction between *Xanthomonas* and *Paenibacillus,* CFUs of *Xanthomonas* and *Paenibacillus* showed a significant strong positive Pearson’s correlation (ϱPearson = 0.87; p-value < 0.001) between resulting CFUs and increasing concentrations of TSB (Fig. 4e), indicating that these two species followed each other’s growth when co-cultured as part of the four-species community. In contrast, neither counts of *Xanthomonas* nor *Paenibacillus* showed a correlation to counts of *Stenotrophomonas* (Fig. S16). To further emphasise the relationship between *Xanthomonas* and *Paenibacillus,* the growth of these two species showed a strong positive significant Pearson’s correlations with each other across time points in the four-species community when cultured in full strength TSB and 50 % TSB (Fig. 4f). This kind of relationship, with a co-increase of counts across time and across different strengths of TSB, was unique for *Xanthomonas* and *Paenibacillus* and could not be identified for any other combination of species in the four-species community (Fig. S17). For 20% TSB, the positive interaction between these two species was not detected and higher CFUs of *Xanthomonas* lead to decreased *Paenibacillus* counts over time.

Of all the four species, *Paenibacillus* responded most strongly to pH alterations in the cultures, with a positive correlation between pH and CFUs for full strength TSB and 50% TSB. In additional support of the pH trend for *Paenibacillus,* counts of CFUs in LB and M9 decreased with increasing pH (for pH > 7.5-8) (Fig. S13), emphasising that the growth of *Paenibacillus* was tightly linked to pH.

### Bacterial induced pH drift in soil

Currently, the pH mediated interaction has only been presented for *in vitro* systems, and as such is only speculative for *in vivo* settings e.g. the rhizosphere associated biofilm communities. To address whether the four species could cause pH alternations in the more natural like systems, pH drive was investigated in bulk soil inoculated with high concentrations of either of the single-species or the four-species cultures. The individual species and the four-species community were inoculated in 5 g of sieved soil with a cell loading of 10^9^ cells per gram of soil. Samples were incubated for eight days, with a vortex induced re-distribution of the soil every second day, before pH was measured in bulk soil (Fig. 5a). All four bacterial species and the four-species consortium were found to significantly increase the pH in bulk soil after eight days of incubation, with *Xanthomonas* promoting the highest increase relative to the control without added bacteria. Plate-spreading of the samples allowed a visual verification of the presence of all four single species in their respective soil samples by colony morphology (data not shown). Similarly *Xanthomonas, Stenotrophomonas* and *Microbacterium* could be identified from the plated soil sample inoculated with the four-species consortium (data not shown).

**Figure 5:**
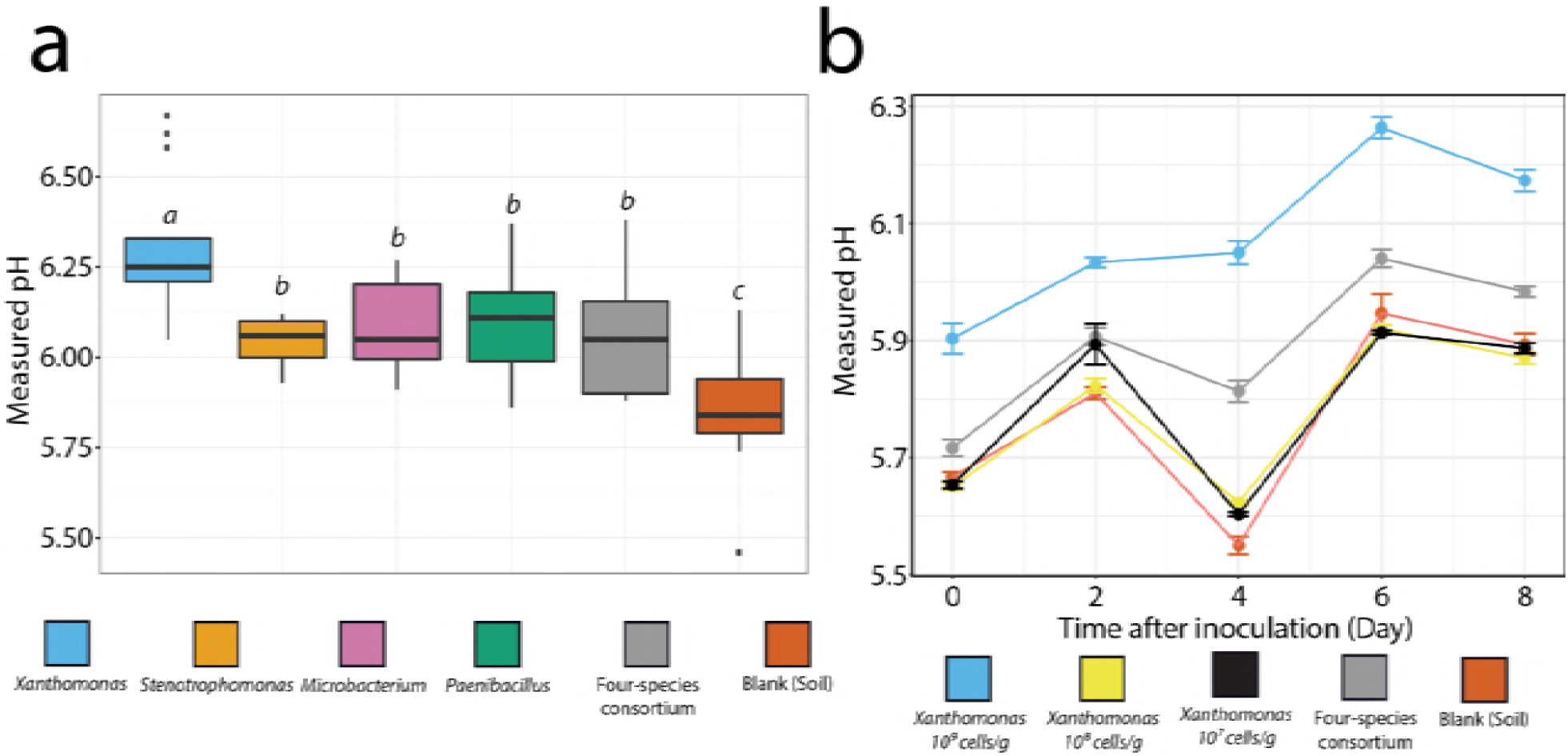
Species mediated pH drift in sieved soil. a) Sieved soil was inoculated with 10^9^ cells per gram of soil with sampling eight days after inoculation. The single species and four-species consortia significantly increased the pH in the soil samples. Statistical grouping of CFU counts with dissimilar letters indicating significant differences with p<0.05 (GLM). Both single- and four-species inoculum promoted a significant pH drift, as compared to blank soil. (n = 5 biological replicates). b) pH over time in soil inoculated with different cell loadings, no bacteria or with the four-species community. (n = 3 biological replicates). High cell loadings were required for pH drift to occur.

Following the pH drift over time in the four-species community and in mono-species cultures of *Xanthomonas* with different cell loadings, showed that most of the pH drift occurred on the first day after inoculation (Fig. 5b). High cell-loadings were required for the effect to occur in bulk soil, as cell loadings below 10^9^ cells per gram of *Xanthomonas* did not provide a significantly increased pH at day 8. Selective CFU counts of *Xanthomonas* were acquired over time to follow the *Xanthomonas* population in mono-species inoculations in the soil or when inoculated into the soil as part of the four-species community. Counts showed that the *Xanthomonas* population was stable from day 0 to 2, and thereafter declined (Fig. S18). Inoculation of either *Xanthomonas* (10^9^ cells per gram) and or the four-species community (with a total of 10^9^ cells per gram) in autoclaved soil also yielded an increased pH in bulk soil after eight days of incubation (Fig. S19). *Xanthomonas* could be selectively isolated from the autoclaved soil samples after inoculation and over time (data not shown).

## Discussion

In the present study, we explored the potential drivers behind a previously observed synergistic interaction between four co-isolated soil bacteria. Opposite pH drive between key community members was found to stabilise the pH environment promoting enhanced growth of selected community members. This mechanisms is very much in line with the type of stabilization presented by Ratzke and Gore (2018) (14). However, other mechanisms besides pH stabilisation could also be in play for the synergy in full strength TSB, as the four-species community had higher total cell counts than the dual-species combination of *Xanthomonas* and *Paenibacillus* where pH stabilisation was also observed. The low cell counts of *Microbacterium* and *Stenotrophomonas* in the four-species community are expected to only cause a negligible pH drive, and the presence of these two isolates might have additional functions in the community. In support, Liu et al. 2017 (32) have shown that inclusion of the *Microbacterium* caused a unique spatial structuring in the four-species community when grown as biofilms under continuous flow. Additionally, cross-feeding on specific amino acids has been suggested/identified as driver for this community in earlier studies (33, 39).

Application of a custom-built x-y-z motorized micro-manipulator setup fixed with micro-sensors enabled us to easily elucidate the interaction of key members in the community by simply addressing a visual interaction on agar plates. Application of micro-sensors to study chemical gradients is a long established technique which has seen diverse application, e.g. within soil sediments (40) or microbial encapsulation in alginate beads (41, 42). Agar plates are routinely used to screen for bacterial interactions and with the diverse range of bacteria which cause pH drift in standard laboratory media (15) one should remember to evaluate the likelihood of pH mediated interaction. Future efforts could apply supporting techniques directly on the agar plates to identify the metabolites causing the interaction, by e.g. utilizing imaging mass spectrometry (43, 44) or chemical imaging (45, 46).

Cellular pH homeostasis is crucial for maintaining functional cells, as intra-cellular proteins function optimally within distinct pH ranges, and because the proton motive force is crucial for bacterial respiration (47, 48). Thus, pH stress can lead to reduced or impaired growth due to defective proteins, a disrupted membrane potential, or the energy cost of maintaining pH homeostasis (47, 49, 50). Changes caused in the pH environment through bacterial growth and its effect on growth of co-cultured bacteria is a well-known fact, with one of the best examples being the syntrophic relationship of *Lactobacillus bulgaricus* and *Streptococcus thermophiles* during yoghurt production (26–29). Hence, that bacteria affect each other by altering the pH environment through their metabolism is not a surprise. Nevertheless, the formulation and predictability of pH drive as a type of inter-species interaction in co-cultures seems not properly established until the recent presentation by Ratzke and Gore (2018) (14). Whether this type of interaction is relevant for natural settings needs to be further established as Ratzke and Gore (2018) performed *in vitro* studies with selected lab isolates with known pH drive and pH growth optimum.

Unlike the observations presented by Ratzke and Gore (2018) (14), our isolates did not undergo ecological suicide when tested as single species, but mono-cultures of *Xanthomonas* and *Paenibacillus* contained lower cell counts compared to those of co-cultures of e.g. *Xanthomonas* and *Paenibacillus* or the four-species community. We found that the pH related interaction was highly media specific and only occurred in high nutrient concentrations. By example, in high media concentration (full strength TSB) both *Xanthomonas* and *Paenibacillus* benefitted from co-cultivation with partners with opposite pH drive (Fig 3.) With medium strength TSB only *Paenbacillus* significantly benefitted from co-cultivation with members with opposite pH drive, e.g. *Stenotrophomonas* or as part of the four-species community (Fig. 4). Hence, with decreasing media strength the interaction faded, suggesting that this type of positive interaction occurs when nutrients is not a limiting factor. When nutrient concentrations are lowered, competition for nutrients becomes a stronger driver in the community, than the positive impact from e.g. pH stabilisation. Notably, this type of interaction might also be stronger in structured systems, such as microbial biofilms, as cooperation has been noted as being stronger in structured environments (17) and cooperating biofilm members tend to evenly mix or co-localise (32, 51, 52). In support, previous studies on biofilms have shown that steep pH gradients can occur within (53) and on the outside (54) of biofilms, generating suitable micro-niches for a diverse set of community members (55).

In perspective to the observations by Ratzke and Gore (2018) (14), and hinting towards the relevance of this type of interactions in natural systems, we observed that pH stabilisation was at least part of the driver behind a previously observed community synergy between our four co-isolated species which are known to form biofilm. As these species were co-isolated from the same decomposing leaf (56), it is likely that they could also occur together in nature and might be able to favour each other’s growth through pH stabilisation in microenvironments under the right conditions. In further support, we observed a pH drive in bulk soil when inoculated with high concentrations of cells, which indicate that i) these bacteria can utilize the nutrients in soil to cause pH drift and ii) a strong pH drift occur in the immediate vicinity (the local micro-environment) of the bacteria in the soil as microbial growth will be centralised around aggregates of nutrients in the soil. Hence, it can be speculated that pH stabilisation might act as a driver for community growth in natural systems, where co-localisation of members creating suitable pH niche for growth can enhance ones fitness in the community.

## Experimental procedures

### Bacterial cultures and strains

The investigated four-species model community was composed of *Stenotrophomonas rhizophila*, *Xanthomonas retroflexus*, *Microbacterium oxydans* and *Paenibacillus amylolyticus*. These isolates were identified during a previous study on plasmid transfer among soil isolates(56) and were later found to exhibit synergistic biofilm formation (31). Bacterial isolates were stored as glycerol stocks at −80°C. From the stocks, the bacterial isolates were streaked onto agar plates containing 1.5% agar-agar (VWR) and 30 g L^−1^ tryptic soy broth (VWR) (TSA). Plates were incubated for 48 hrs at 24°C. Single colonies were used to inoculate 5 mL tryptic soy broth, 30 g L^−1^ tryptic soy broth (VWR) (TSB). 5 mL cultures were incubated over night at 250 rpm at 24°C.

### Cultivation in 24 well plates

For experiments with full strength TSB, over-night bacterial cultures were directly diluted to an optical density at 600 nm (OD_600_) of 0.15 before use in either TSB or, where noted, in TSB complemented with 5 mM nitrate. For testing the effect of media composition, diluted variants of TSB were included along with LB broth (LB) and a mixed minimal medium (M9). LB (25 g L^−1^; LB broth (Miller); VWR) was included due to its complexity, to address if the pH synergy would occur in other complex media types. M9 (10.5 g L^−1^; M9 broth; Sigma-Aldrich) was complemented with 0.5% (w/v) tryptone (Tryptone enzymatic digest from casein; Sigma-Aldrich) and 0.5% (w/v) glucose (D(+)-Glucose; Merck) as the nitrogen and carbon sources to have a limited media. For preparation of cell cultures for the different media variants, cells from over-night cultures were precipitated by centrifugation at 5000g for 5 min. The supernatant was discarded, and cells were washed in 0.9% NaCl (w/v) before re-dissolving the cells. Cells were re-precipitated by centrifugation and the supernatant was discarded before the cell were re-dissolved in the appropriated media. Cultures were then adjusted to an OD_600_ of 0.15 before use. OD_600_ adjusted cell cultures were used to inoculate 24 well plates with mono- and co-cultures. All wells contained a total of 2 mL of OD_600_ adjusted culture. For mono-species cultures, 2 mL of the single species culture was used, while equal volumes of each species were used for co-cultures. Inoculated plates were incubated under static conditions for up to 48 hrs at 24°C.

For CFU counts from 24 well plates, cultures were homogenized with a pipette and diluted in 1xPBS. Diluted culture was plated on TSA (15 g L^−1^ agar powder (VWR) and 30 g L^−1^tryptic soy broth (Sigma-Aldrich)) plates complemented with 40 µg mL^−1^ congo red (Fluka), and 20 µg mL^−1^ Coomassie brilliant blue G250 (Sigma-Aldrich). Colony forming units was counted after 48 hrs of incubation at 24°C by differentiating species based on dissimilar colony morphology.

### Agar plates

All experiments, including pH and O_2_ measurements, performed on agar plate colonies was performed on 50% TSA plates (15 g L^−1^ agar powder (VWR) and 15 g L^−1^ tryptic soy broth (Sigma-Aldrich)). Plates for visualization of morphological changes were 50% TSA plates complemented with 40 μg mL^−1^ congo red (Fluka) and 20 μg mL^−1^ coomassie brilliant blue G250 (Sigma-Aldrich), referred to as Congo Red plates.

Colony spotting for pH and O_2_ measurements was done with a fixed distance between colony centers. The spotting (interaction zone) area was divided into a grid, with each grid-square being 2.5 × 2.5 mm. Colonies were spotted with an approximate distance of 1.25 cm between colony centers. 5 μL of OD_600_ 0.15 adjusted cultures (prepared as previously described) were used for spotting bacterial colonies. Similarly, two-species interaction studies were performed with an approximate distance of 1.25 cm between the center of the colonies. 5 μL of OD_600_ 0.15 adjusted cultures (prepared as previously described) were spotted.

Buffer stabilized plates were 50% TSA agar plates complemented with 200 mM sodium-acetate (pH 5 and 5.5), potassium-phosphate (pH 6, 6.5 and 7), Trisma base (pH 7.5, 8, 8.5 and 9), and sodium-carbonate (pH 9.5 and 10.5) buffers.

### Data analysis and plotting

Boxplots were plotted using the ggplot2 R package. For boxplot with CFU and pH, box width was set to 2x the standard error of the measured pH within each group. Statistical significance was inferred between groups e.g. on log2(CFU counts) with a generalised linear model with Tukey pairwise comparison and multiple hypothesis testing by Single step method using the multcomp package in the R environment (referred to as GLM) (57). Spearman’s ranked correlations were used to infer correlations between CFU and endpoint pH, and Pearson’s correlations was used to infer correlations between CFU counts of two species.

### Soil Samples

Soil from a Danish research field (Taastrup, Denmark, Coordinates; 55.669762, 12.300498) was sieved for particles < 2mm and the soil was stored cold until use. This soil was chosen as the bacterial isolates were originally isolated from soil obtained from the same research facility. Soil samples contained 5g soil, contained in 50 mL Falcon tubes. The soil was inoculated with 2 mL of bacterial culture with varying inoculation sizes of bacteria. Cells from over-night bacterial cultures in TSB were precipitated by centrifugation at 5000g for 5 min and the supernatant was discarded. Cells were washed in phosphate buffered saline (1xPBS) and re-precipitated by centrifugation. Cells were re-dissolved in 2mL 1xPBS and adjusted to the appropriate OD600 to provide 2 mL cell suspension with cells to yield e.g. 10^9^ cell per gram of soil. For mix cultures cells were mixed in equal proportions to yield a total of e.g. 10^9^ cell per gram of soil. Cell suspensions were used to inoculate the 5 g soil samples. The addition of 2 mL solution left the soil with a very thin water film on top of the soil. Samples were vortexed for 5 sec. to distribute liquid and bacteria in the soil. Samples were incubated at 24°C under static conditions. On every second day, the tubes were briefly vortexed to re-distribute nutrients and cells in the soil. Blank samples without inoculation of bacteria were prepared by inoculating the soil with 2 mL 1xPBS. For sampling 5 mL sterile water was added to the tubes, and the tubes were shaken for 10 min. before pH was measured in the water fraction of the sample. To verify the presence of the inoculated species in the soil 100 μL of the water suspension was serial diluted in 1xPBS and plate-spread on TSA plates complemented with Congo red and Coomassie brilliant blue G250, as described for 24 well plates. Inoculated species could be recognized by their unique colony morphology. For selective counts of *Xanthomonas*, agar plates were further complemented with 20 μg mL^−1^ Kanamycin.

### Microsensor measurements

2D microsensor measurements of pH and O_2_ concentration transects across agar plates were conducted with the microsensors mounted in a custom-built x-y-z motorized micromanipulator setup fixed to a heavy stand (58). Similar motorized x-y-z micromanipulator setups can be obtained from commercial sources; e.g. *Pyro-Science GmbH, Aachen, Germany or Unisense A/S, Aarhus, Denmark.*

For O_2_ measurements, a fiber-optic O_2_ microsensor (OXR50-HS, tip diameter 50 μm) was connected to an O_2_ meter (FireStingO2); both components were obtained from Pyro-Science GmbH Aachen, Germany (pyro-science.com). Calibration of the microsensor was performed as specified by the manufacturer by measurements in air saturated and O_2_ free water, respectively.

For pH measurements, we used a pH glass microelectrode (tip diameter 50 μm, pH50; Unisense A/S) in combination with a reference electrode (tip diameter of ~5 mm; Unisense A/S) immersed in the agar plate. Both sensors were connected to a high impedance pH/mV-Meter (Unisense A/S). Before measurements commenced, the pH microelectrode was linearly calibrated from sensor mV readings in three pre-known pH buffers (pH 4, 7 and 9) showing a log-linear response to [H^+^] of ~51 mV/pH unit at experimental temperature (24°C ± 0.5°C).

For N_2_O measurements, a microsensor (tip diameter 50 µm, N2O50; Unisense A/S) was connected to a PA2000 pico-amperometer (discontinued product from Unisense A/S). The sensor was pre-activated, polarized and calibrated as stated in the manual using sensor readings in N_2_O free water and then after addition of known amounts of N_2_O saturated water.

A USB microscope (dino-lite.eu, model AM7515MZTL) was used to determine when the microsensor tip touched the surface of the agar plate. All 2D measurements (pH and O_2_) were conducted at a depth of ≈100 µm below the surface. A custom-made profiling software (Volfix; programmed by Roland Thar) was used to control the *x-y-z* motorized micromanipulator and to read out both sensor signals. A similar software, Profix, can be downloaded free of charge from pyro-science.com. An analog to digital converter (ADC-101; Pico Technology, UK) had to be used in order to interface the profiling software with the O2 meter (using the analog output of the FireStingO2) and the pH/mV-Meter. Time course measurements, of e.g. O2, in static culture were recorded with free logging software (SensorTrace logger; Unisense A/S).

## Acknowledgement

We thank Annette Løth for assistance with media preparation, Maja Holm Wahlgren and Professor Anders Primé for their assistance with nitrous oxide measurements, and Peter Østrup Jensen for assistance with microsensor equipment. We thank Esben Nielsen and Gosha Sylvester for their assistance with ammonium, and nitrate/nitrite measurements. Lastly, we thank Jakob Russel for assistance with data handling in the R environment.

